# Systematic Multi-Omics Investigation of Androgen Receptor Driven Gene Expression and Epigenetics changes in Prostate Cancer

**DOI:** 10.1101/2024.07.22.604505

**Authors:** Lin Li, Kyung Hyun Cho, Xiuping Yu, Siyuan Cheng

**Affiliations:** Department of Biochemistry and Molecular biology, LSU Health Shreveport, Shreveport, LA; Feist-Weiller Cancer Center, LSU Health Shreveport, Shreveport, LA; Department of Urology, LSU Health Shreveport, Shreveport, LA

## Abstract

**Background:** Prostate cancer, a common malignancy, is driven by androgen receptor (AR) signaling. Understanding the function of AR signaling is critical for prostate cancer research. **Methods:** We performed multi-omics data analysis for the AR^+^, androgen-sensitive LNCaP cell line, focusing on gene expression (RNAseq), chromatin accessibility (ATACseq), and transcription factor binding (ChIPseq). High-quality datasets were curated from public repositories and processed using state-of-the-art bioinformatics tools. **Results:** Our analysis identified 1004 up-regulated and 707 down-regulated genes in response to androgen deprivation therapy (ADT) which diminished AR signaling activity. Gene-set enrichment analysis revealed that AR signaling influences pathways related to neuron differentiation, cell adhesion, P53 signaling, and inflammation. ATACseq and ChIPseq data demonstrated that as a transcription factor, AR primarily binds to distal enhancers, influencing chromatin modifications without affecting proximal promoter regions. In addition, the AR-induced genes maintained higher active chromatin states than AR-inhibited genes, even under ADT conditions. Furthermore, ADT did not directly induce neuroendocrine differentiation in LNCaP cells, suggesting a complex mechanism behind neuroendocrine prostate cancer development. In addition, a publicly available online application LNCaP-ADT (https://pcatools.shinyapps.io/shinyADT/) was launched for users to visualize and browse data generated by this study. **Conclusion:** This study provides a comprehensive multi-omics dataset, elucidating the role of AR signaling in prostate cancer at the transcriptomic and epigenomic levels. The reprocessed data is publicly available, offering a valuable resource for future prostate cancer research.

## 1. Introduction

Prostate cancer is a prevalent malignancy intricately linked to androgen receptor (AR) signaling, with most cases being androgen-sensitive and dependent on AR’s transcriptional activity^1^. AR signaling begins when androgens bind to AR, causing its translocation from the cytosol to the nucleus, where it regulates target gene expression^2^. This process involves the displacement of heat shock proteins (HSPs), triggering AR dimerization, phosphorylation, and the necessary conformational changes^3^.

AR acts as a central mediator in prostate cancer, providing growth signals that are crucial for the cancer cells’ development and progression^4–6^. Understanding the AR cistrome, the complete set of AR genomic binding sites, is essential for prostate cancer research, especially for pathogenesis, treatment, and progression^7–10^. Key AR-regulated genes, such as PSA (KLK3), KLK2, and FKBP5, and transcriptional regulators like p300, CBP, HDACs, and LSD1, have been extensively characterized^11–13^. Interestingly, researchers found that unlike other transcription factors binding to proximal promoter regions of regulated genes, AR binding sites are often far from (>10 kb) the transcription start sites (TSS) of regulated genes^14^. Subsequent studies identified that AR directly upregulates canonical androgen-responsive genes like PSA and TMPRSS2 through chromatin looping interactions, highlighting chromatin looping as a crucial mechanism that facilitates interactions between distal enhancers and proximal promoters^15–17^. Furthermore, the AR cistrome is tissue specific. Key contributors to AR cistrome reprogramming in prostate cancer cells include Forkhead Box A1 (FOXA1), homeobox protein HOXB13, GATA binding protein 2 (GATA2), and ETS-related gene (ERG)^18–20^.

Androgen deprivation therapy (ADT), which reduces testosterone levels and blocks AR, is a cornerstone in treating advanced prostate cancer, particularly when surgical or radiation options have been exhausted^21,22^. Charcoal-stripped fetal bovine serum (CS-FBS) and Enzalutamide (ENZ), a second-generation AR antagonist, are frequently used to study androgen responsiveness in cultured prostate cancer cells^23–25^. Switching prostate cancer cells from FBS to CS-FBS reduced androgen receptor activity and inhibited cell proliferation^23,26^. Additionally, treating LNCaP cells with enzalutamide sensitized them to radiotherapy, enhanced apoptosis, delayed DNA repair, and decreased colony survival^27,28^.

The development of high-throughput sequencing techniques has revolutionized prostate cancer research by significantly increasing the number of readouts following treatments like CS-FBS or Enzalutamide. For example, RNA sequencing (RNAseq) provides comprehensive insights into gene expression profiles, enabling the identification of key genes and pathways involved in prostate cancer progression and treatment response. RNAseq has been instrumental in characterizing gene expression changes in prostate cancer cell lines under various conditions, such as androgen/anti-androgen treatment^29^. The Assay for Transposase-Accessible Chromatin using sequencing (ATACseq) allows researchers to investigate chromatin accessibility, providing insights into open chromatin regions and transcription factor binding sites^30^. Chromatin immunoprecipitation followed by high-throughput sequencing (ChIPseq) is a powerful tool to describe chromatin modification landscapes and transcription factor binding sites. Several studies have utilized ChIPseq to map AR binding sites in prostate cancer cell lines, providing valuable insights into prostate cancer progression mechanisms^14,20^.

The Gene Expression Omnibus (GEO) database serves as a commonly used public repository for high-throughput sequencing data, including RNAseq, ATACseq, and ChIPseq^31^. Re-analyzing and curating these multi-omics data offers a valuable opportunity to gain comprehensive and unbiased insights into gene regulatory mechanisms in prostate cancer research. In this study, we provide the bioinformatics analysis of AR signaling in terms of gene expression, chromatin accessibility, cistrome, and transcription factor co-occupancy using multi-omics data derived from a single AR^+^, androgen-sensitive LNCaP cell line^32^. The use of a single cell line, LNCaP, allows us to eliminate the effects of different genetic backgrounds and cellular contexts on AR signaling, ensuring that our findings are specifically attributed to the AR pathway itself. Furthermore, the unbiased bioinformatics analysis performed in this study allows for a more objective description of AR signaling. This approach highlights AR’s central role and its regulatory network in managing prostate cancer and underscores the significance of high-throughput sequencing technologies in advancing our understanding of these mechanisms. Also, this the high-quality re-processed multi-omics data were deposited to the public cohort that can be easily reused for other research projects.

## 2. Result

### 2.1 Multi-omics data collection and processing

To establish the multi-omics dataset for the prostate cancer LNCaP cell line, data was collected and re-processed as detailed in the methods section. In short, the raw files and metadata were acquired from the SRA and GEO databases. After quality control, data of 215 RNAseq from 52 different studies, 144 ATACseq from 24 studies, and 359 ChIPseq from 52 studies were included in the downstream analysis. The RNAseq samples were further classified into 3 groups: control (no treatment or control treatments), charcoal-stripped fetal bovine serum treated (CS-FBS) and Enzalutamide (ENZ) treated. The ATACseq and ChIPseq samples were labeled by simplified treatment information and the non-treated (NT), dihydrotestosterone (DHT)/R1881 treated (androgen^+^) and CS-FBS/Bicalutamide/Enzalutamide treated (androgen^-^). Although only ADT related samples were included in the analysis below, all processed data were deposited into Figshare and open to download. The individual links were listed in data availability section.

### 2.2 Transcriptome analysis identified AR regulated genes and functions

The transcriptome identity of each RNAseq sample was described using principal component analysis (PCA) of the TPM value of each sample. As shown in Figure 1A, the samples from the CS-FBS and ENZ groups located together while separating from the control group, indicating that androgen-targeted treatments (CS-FBS & ENZ) had a similar effect on the LNCaP cell transcriptome. The PCA result also suggested this treatment effect size was substantial enough for downstream differential gene expression analysis.

**Figure 1.**
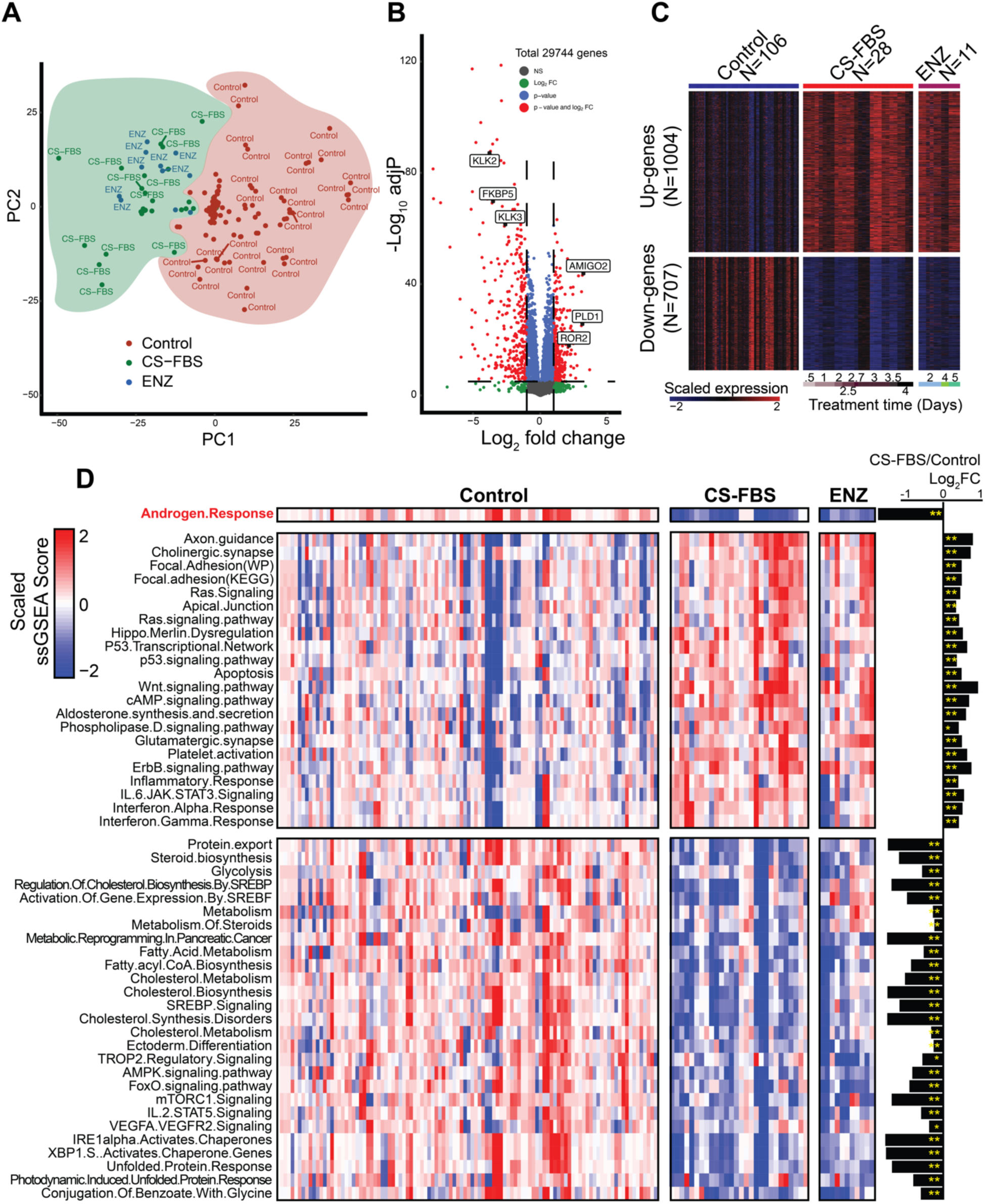
Transcriptomic Analysis of LNCaP Cells under Androgen Targeted Treatments. (A) Principal Component Analysis (PCA) of RNAseq samples based on TPM values. Samples from CS-FBS (blue) and ENZ (green) groups cluster together, separated from Control (red) samples. (B) Volcano plot displaying differentially expressed genes between CS-FBS and Control groups. Up-regulated (red) and down-regulated (blue) genes are identified. (C) Heatmap visualizing the expression of the identified up-regulated and down-regulated genes across Control, CS-FBS, and ENZ samples. (D) Heatmap of single-sample gene-set enrichment analysis (ssGSEA) scores for various gene sets, including “androgen response”. The right part shows a bar plot of the log_2_ fold-change of each gene-set activity from CS-FBS compared to Control group samples. The asterisks (*) indicate t-test p-values less than 0.05, and double asterisks (**) indicate p-values less than 0.01.

To identify AR-regulated genes and describe AR signaling function in LNCaP cells, we identified 1004 ADT up-regulated genes and 707 down-regulated genes by comparing the transcriptomes of CS-FBS group samples with control group samples (Figure 1B). The ENZ group samples were not considered due to the smaller sample number. As expected, known AR-activated genes (KLK2, KLK3, FKBP5) were significantly decreased in CS-FBS samples, indicating the success of this transcriptome study (Figure 1B). Then the up- and down-regulated genes were visualized by a heatmap (Figure 1C). Notably, ENZ group samples showed a similar gene expression trend to CS-FBS samples, although they were not included in the DE analysis. Furthermore, these DE genes showed no significant correlation with ADT treatment time, as annotated at the bottom of the heatmap. This result suggested the response of AR signaling to treatment is rapid and stable.

To further explore the biological significance of the AR-regulated genes, we used a combination of gene-set enrichment and single-sample gene-set enrichment (ssGSEA). First, the official gene symbols of up- and down-regulated genes were enriched against the gene-set library from “Enrichr”^33^. Then, significant gene sets were collected and scored for each RNAseq sample using the ssGSEA method to increase analysis resolution and ensure global gene coverage within each sample. As shown in Figure 1D, the “androgen response” activities were high in control samples but lower in CS-FBS and ENZ samples. Additionally, the different levels of “androgen response” activities within each group explained the previously noted varying expression patterns of DE genes within the same group (Figure 1C). Since the sample order (columns of the heatmap) was kept the same between gene expression and gene-set activity heatmaps (Figure 1C&D), combining both heatmaps showed that the varying expression levels of the DE genes was associated with different calculated AR signaling activity which was contributed by different treatment strategies and doses. Taking together, in LNCaP cells, the AR signaling responses to treatment in a rapid, stable and dose dependent manner.

Besides androgen-response, the other enriched gene-sets revealed the downstream functions of AR signaling. Three neuron-related pathways (axon guidance, cholinergic synapse, and glutamatergic synapse) were enriched in CS-FBS and ENZ samples, indicating that AR signaling inhibits neuron-related gene expression. Under ADT, the expression of these neuron-related genes may contribute to the partial neuronal features of LNCaP cells, particularly the previously reported neuronal-like (spindle-like) morphology^34,35^. Additionally, the increase in cell-cell contact pathways (focal adhesion and apical junction) may be associated with these morphological changes. Besides, the activation of the P53 signaling pathway and apoptosis-related genes may partially explain the decreased cell proliferation in LNCaP cells under ADT.

Tumorigenic signaling pathways, such as Wnt, Hippo, and ErbB, were activated in ADT samples, suggesting an inhibitory role of AR on these pathways. These findings supported the dual function of AR signaling in prostate cancer: both oncogenic and tumor-suppressive^36,37^. Interestingly, inflammation-related signaling pathways were inhibited by AR and elevated in ADT samples as recently reported and consistent with the critical role of inflammatory signaling pathways during the development of castration-resistant prostate cancer (CRPCa) ^38–40^.

While the ADT induced gene-sets that shed light on potential CRPCa mechanisms, the ADT inactivated gene-sets can directly describe the function of AR signaling in AdPCa cells. Consistent with previous reports, AR regulates protein export and metabolism in LNCaP cells^41^. Besides, similar to our data that AR inhibits neuronal-related gene expression, the decreased ectoderm-differentiation gene-set score in ADT samples also suggests AR’s role in maintaining and promoting the prostatic differentiation of LNCaP cells.

### 2.3 Androgen deprivation won’t affect proximal promoter chromatin status of AR-regulated genes

Besides the AR downstream genes and their functions revealed by transcriptome study, the ATACseq and ChIPseq studies can further describe the function of AR in epigenetics modifications. Based on previous transcriptome study, all human GRCh38 protein-coding genes were categorized into four groups: (1) genes up-regulated upon ADT (n=1004); (2) genes down-regulated upon ADT(n=707); (3) consistently highly expressed genes (average TPM >3 with no significant expression change upon ADT); and (4) consistently low expressed genes. Consequently, the genomic regions of these genes were used for visualizations.

ATAC-seq, which depicts accessible regulatory regions in the genome, revealed that proximal regulatory regions (3kb upstream and downstream of the transcription start site) were not affected by androgen status, even in AR-regulated gene regions. As shown in Figure 2A, chromatin open regions (indicated by ATACseq peaks) were centered around the transcription start site (TSS) in consistently high expression gene regions but not in consistently low expression regions, indicated the correlation between promoter chromatin accessibility and gene expression^42^. While comparing the Androgen^+^ and Androgen^-^ samples, both the ADT up- and down-regulated gene regions exhibited almost identical ATACseq peaks in 2 groups of samples, as evidenced by the heatmap, which shows individual genomic regions, and the profile plot, which displays the average ATACseq signal intensities (Figure 2A). This result suggested that ADT did not alter the global chromatin accessibility patterns and, more surprisingly, did not affect the accessibility of AR-regulated gene regions.

**Figure 2.**
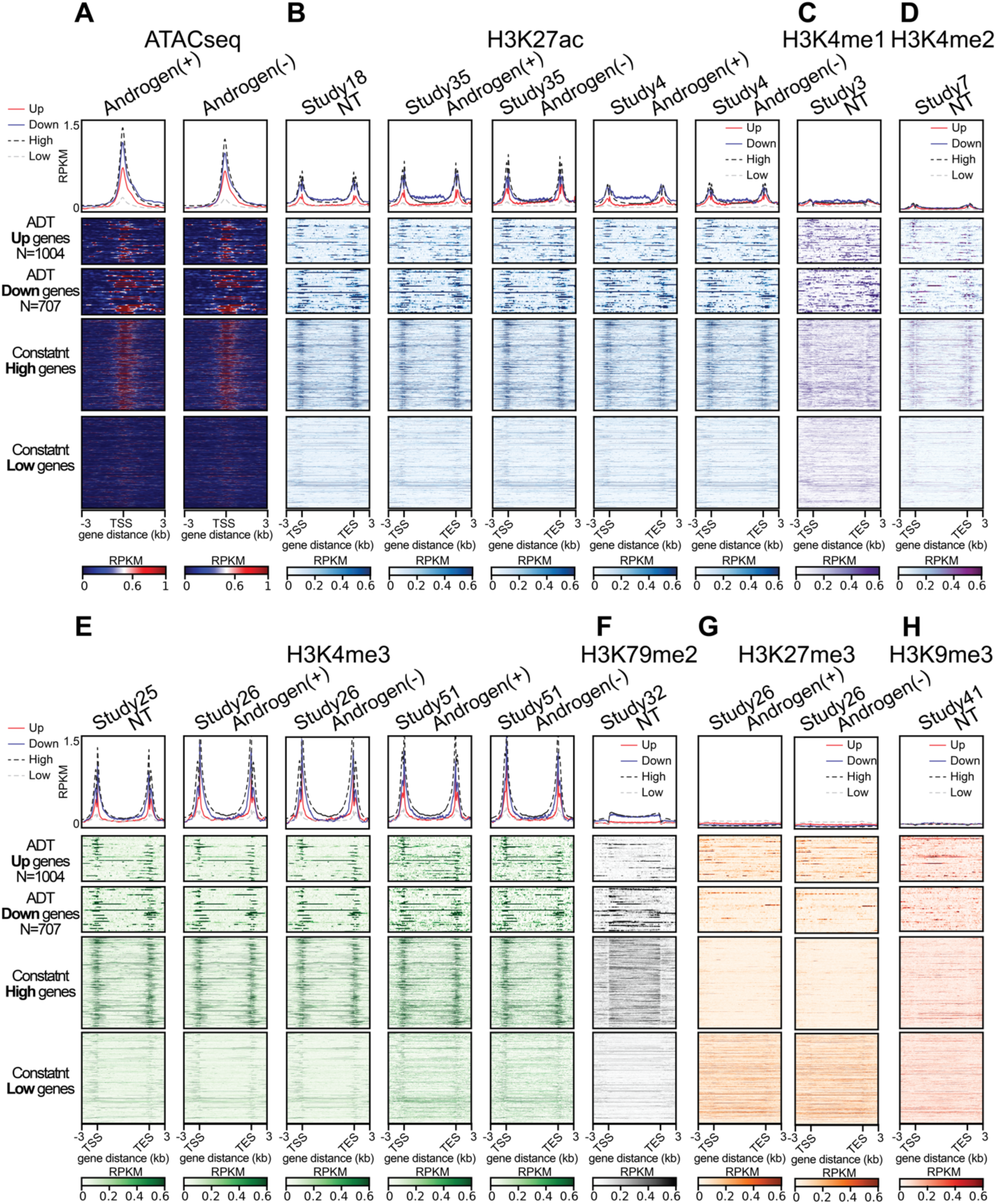
Chromatin Accessibility and Histone Modification Profiles in ADT Regulated Genes Locus. (A) ATAC-seq analysis showing chromatin accessibility around transcription start sites (TSS) in androgen-positive (+) and androgen-negative (-) conditions. Heatmaps display reads per kilobase per million mapped reads (RPKM) values for ADT up-regulated genes (N=1004), ADT down-regulated genes (N=707), and genes with consistently high or low expression. ChIP-seq analysis of various histone modifications including H3K27ac (B), H3K4me1 (C), H3K4me2 (D), H3K4me3 (E), H3K79me2 (F), H3K27me3 (G) and K3K9me3 (H) under androgen-positive and androgen-negative conditions. Heatmaps show RPKM values across the same gene regions as panel A. Profile plots above each heatmap represent average RPKM values across the regions of interest, centered on the TSS and extending ±3 kb.

In addition to ATACseq, ChIPseq targeting various chromatin modifications provided further insights. H3K27ac (histone H3 lysine 27 acetylation) marks active enhancers and promoters, while H3K4me3 (histone H3 lysine 4 trimethylation) marks active promoters^43,44^. H3K4me1 (histone H3 lysine 4 monomethylation) marks both active and poised enhancers^45,46^, and H3K4me2 (histone H3 lysine 4 dimethylation) is known as a signature for predicting enhancers^47^. As shown in Figure 2B-E, all three groups of genomic regions (up, down, and high) had high levels of H3K27ac and H3K4me3, and low levels of H3K4me1/2, indicating that these regions are dominantly active promoters. This is consistent with the well-known fact that promoters are usually located near transcription start sites (TSS)^48^. In contrast, the silenced gene markers H3K27me3 (histone H3 lysine 27 trimethylation) and H3K9me3 (histone H3 lysine 9 trimethylation) were relatively lower in these 3 groups of regions but higher in constant-low regions (Figure 2G-H).

Comparing these ChIPseq samples under different androgen statuses revealed results similar to those from ATACseq. In these proximal promoter regions, ADT in LNCaP cells did not affect chromatin modifications, including H3K27ac (Figure 2B) and H3K4me3 (Figure 3E), as depicted by the almost identical heatmaps and profile plots. In summary, although AR signaling significantly affects its downstream genes expression, it does not modify chromatin in the proximal promoter regions of these genes.

**Figure 3.**
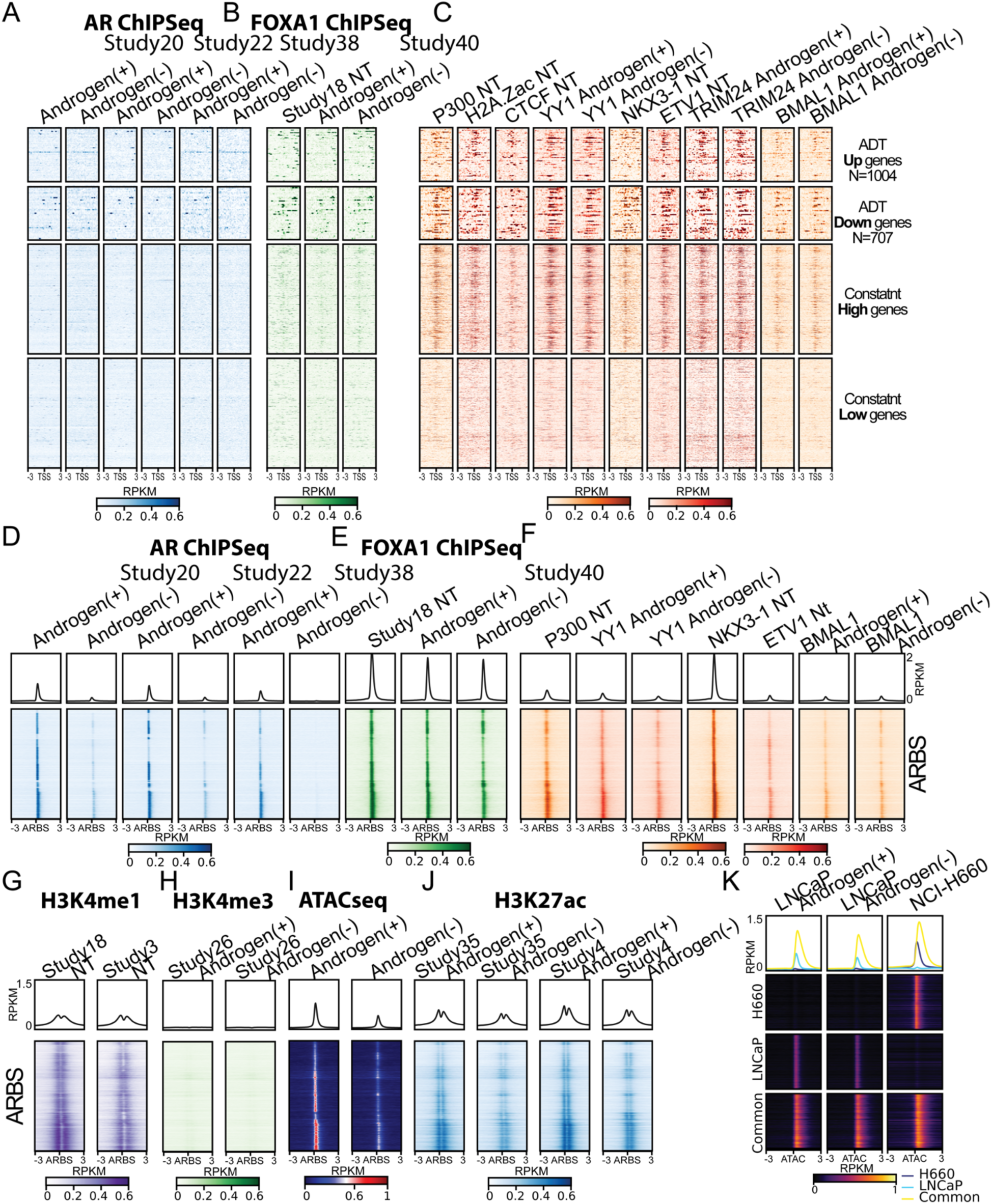
Chromatin accessibility, histone modification, and transcription factors binding profile of AR regulated gene locus and AR binding sites (ARBS). The transcription factors binding on different gene locus were visualized by heatmap for AR (A), FOXA1 (B) and others (C). These factors binding on ARBS were also visualized by heatmaps and profile plots (D-F). Then the chromatin modifications on ARBS including H3K4me1 (G), H3K4me3 (H), H3K27ac (J) and chromatin accessibility on ARBS revealed by ATACseq were also visualized by heatmaps and profile plots. (K) The ATACseq signals from LNCaP androgen-positive, LNCaP androgen-negative and NEPCa cell line H660 were visualized by heatmaps and profile plots focusing on H660-specific, LNCaP-specific and common ATACseq peaks.

Interestingly, by comparing the ADT up- and down-regulated genes’ region, both ATACseq and ChIPseq data revealed differences. The ADT down-regulated (AR-induced) genes showed higher chromatin accessibility (ATACseq) and maintained higher levels of active chromatin markers (H3K27ac, H3K4me3 ChIPseq) compared to ADT up-regulated (AR-inhibited) genes (Figure 2A-E). Additionally, this difference was not affected by androgen status, suggesting that even without AR signaling activity, the AR-induced genes were still more epigenetically active suggesting that these genes, at least in LNCaP cells, were primed for high-level transcription (Figure 2A-E). The H3K79me2 (histone H3 lysine 79 dimethylation) mark, associated with highly transcribed genes^49^, also partially supported these results. As shown in Figure 2F, under normal culture conditions (non-treated), the AR-induced genes (ADT down-regulated genes) maintained similar levels of this high-expression chromatin mark as the constant-high genes while the AR-inhibited genes did not.

### 2.4 AR binding in proximal promoter regions was absent

While the previous result identified AR has no or marginal effect to the proximal promoter of its downstream genes, the further transcription factor ChIPseq data analysis answered that observation well. As shown in Figure3A, the AR ChIPseq suggested all four groups of the proximal promoter regions had no enrichment of AR binding under both Androgen^+^ and Androgen^-^ conditions. In addition, the binding signal of FOXA1, a prostate essential pioneer transcription factor^50^, wasn’t enriched in these regions (Figure 3B). Unlike AR and FOXA1, another prostate specific transcription factor NKX3-1 was detected in AR induced, constant-high and some AR inhibited gene regions (Figure3C).

Since this study re-processed all available ChIPseq data from LNCaP cells, other transcription factors binding to the promoter regions were also analyzed. First, P300, a histone acetyltransferase, and its downstream H2A.Zac (histone H2A.Z acetylation) were visualized to highlight the TSS of the transcriptionally active gene loci^51,52^ (Figure 3C). Also, previously reported AR-associated transcription factors like YY1^53^, CTCF^54^, ETV1^55,56^, and TRIM24^57^ were detected in the proximal promoter regions, suggesting their role in regulating AR downstream genes (Figure 3C). BMAL1, a transcription factor that regulates the expression of genes associated with lipid metabolism, adipogenesis, and inflammatory responses^58,59^, also showed binding signals in both ADT up- and down-regulated gene regions (Figure 3C). This finding might explain the previously observed enrichment of metabolism and inflammation functions in ADT regulated genes (Figure 1D).

### 2.5 AR bound to distal enhancer and affects chromatin modifications

Although previous research and reviews have already proposed that AR primarily binds to distal regions of regulated genes^60^, our data above suggesting the non-enrichment of AR binding, even in the promoters of AR-induced genes, was still astonishing. We then visualized the re-processed multi-omics data, focusing only on the AR binding sites (ARBS) detected by AR ChIPseq data. As shown in Figure 3D, AR binding signals were consistently detected in three independent studies when androgen was present but diminished with ADT (androgen^-^). Additionally, FOXA1 binding signals were highly detected in these ARBS and were not affected by ADT (Figure 3E). These results further support the known function of FOXA1 in opening chromatin structure for AR binding^18^. Interestingly, NKX3-1 binding was also highly detected in the ARBS, suggesting its co-localization with AR in these regions (Figure 3F).

The H2A.Zac was not detected (H2A.Z was not detected neither, data not shown), indicating the limited presence of TSS in ARBS (Figure 3F). In addition, the chromatin ChIPseq data showed that the ARBS enriched H3K4me1 and H3K27ac, but not H3K4me3, suggesting that ARBS are located at enhancers rather than promoters (Figure 3G&3H). As for the AR-associated transcription factors, the binding of YY1, ETV1, and BMAL1 were all detected in ARBS (Figure 3F). However, TRIM24 binding was not detected in ARBS, suggesting it might not recruit or be recruited by AR (data not shown). The P300 binding was also enriched in the ARBS (Figure 3F).

Additionally, ATACseq showed that the ARBS were less accessible under ADT (androgen^-^) conditions suggested that AR is capable to modify chromatin accessibility across its binding enhancer regions. When AR signaling is inhibited, these enhancers are relatively closed (Figure 3I). Besides, H3K27ac, the active chromatin marker, was also decreased in ADT samples (Figure 3J). Taking these big data together, we further supported and refined the previous theory that AR activates gene transcription by binding to distal enhancers and recruiting P300 for chromatin acetylation^62,63^.

### 2.6 The treatment induced neuroendocrine differentiation is not regulated by AR directly

Clinically, prolonged androgen targeting therapy has been associated with the emergence of the neuroendocrine (NE) phenotype in prostate cancer. Neuroendocrine prostate cancer cells express NE marker genes, lose AR expression, and do not respond to ADT, which contributes to poor prognosis, treatment resistance, and aggressiveness^64^. Unfortunately, the mechanism behind NE phenotype development remains unclear. Recent studies have shown that AR^+^ prostate adenocarcinoma (AdPCa) cells can transform into neuroendocrine prostate cancer (NEPCa) cells through ADT in xenograft models^65,66^. Interestingly, another study suggested that NEPC development does not require prior ADT^67^. For investigation, we analyzed our multi-omics data and provided unbiased insight.

Initially, we examined the effect of ADT on NEPCa development using transcriptome data. We previously identified 169 NEPCa marker genes by overlapping the differentially expressed genes derived from multiple studies^68^. From these genes, only 16 showed a significant increase (adjust p-value < 0.05 & log2 fold-change > 0.5) in CS-FBS samples compared to control samples. The majority of the NEPCa marker genes (153 genes) were either lowly expressed or showed no significant expression changes. All these gene expression data can be accessed through our LNCaP-ADT online website at https://pcatools.shinyapps.io/shinyADT/. Additionally, neither the functional enrichment of differentially expressed genes nor ssGSEA identified NEPCa marker genes as enriched in ADT samples (data not shown).

Next, we characterized the regulatory regions in LNCaP and a NEPCa model cell line, NCI-H660 (H660), using ATACseq data. As shown in Figure 3K, LNCaP and H660 maintained distinct ATACseq peak profiles, indicating their different chromatin accessibility landscapes. In contrast to the differences between LNCaP and H660, AR signaling did not open H660 unique peaks, as shown by the nearly identical heatmaps of LNCaP with and without androgen presence (Figure 3K). On the other hand, ADT samples exhibited slightly lower LNCaP-specific peaks, likely due to decreased ATACseq signals at ARBS (Figures 3G and 3K). In summary, ADT neither directly triggered NE gene expression nor initiated chromatin remodeling towards NEPCa. While if taking together our previous data on ADT downstream genes’ functional enrichment, the role of ADT during NEPCa development may involve releasing the inhibitory effect of AR on the differentiation-related genes in individual cells and promoting the selection of NEPCa cells within the tumor.

## 3. Methods

### 3.1 Data collection, annotation and acquiring

The multi-omics data from the Sequence Read Archive (SRA) were fetched using a pipeline we previously built^69^. Briefly, the “Entrez Direct” toolkit was used to retrieve all sequencing data records from LNCaP cells by searching the keyword “lncap.” The corresponding experiment metadata for the fetched SRA records were accessed using the “GEOquery” R package^70^. After removing mismatched samples, a total of 2,031 multi-omics sequencing samples were included for downstream analysis. Considering the complexity of ChIPseq experiments, only those studies with available loading controls (IgG or input DNA) were included.

The experimental design and detailed information were manually inspected and simplified. For instance, samples from the same experiment were grouped and named sequentially. The “E” in sample names stands for “Experiment” and is followed by a sequential number to distinguish samples from different experiments (each). Treatment information was simplified for better understanding, while the original GEO accession number was retained for each sample for further details. The raw sequencing files were downloaded using the “SRA-Toolkit” and stored in “.fastq.gz” format for downstream analysis.

For better organization, we created the Integrative Multi-Omics Prostate Cancer Informatics Initiative (IMPACT) to collect all multi-omics data we re-processed for LNCaP cells and for other prostate cancer cells in the future.

### 3.2 General data processing

The sequencing quality was assessed using “FastQC” (https://github.com/s-andrews/FastQC) and visualized with “MultiQC” (https://github.com/MultiQC/MultiQC). Subsequently, all samples were trimmed to remove sequencing adaptors and low-quality reads using “Trim Galore” (https://github.com/FelixKrueger/TrimGalore). The latest genome assembly, GRCh38, was utilized as the reference genome for sequencing alignment. The genome file (.fa) and reference file (.gtf), along with other necessary files, were obtained from AWS iGenomes (https://github.com/ewels/AWS-iGenomes). All analyses were conducted in a Linux Ubuntu environment on a local workstation.

### 3.3 RNAseq data processing, analysis and visualization

The “nf-core/rnaseq” pipeline was used for processing RNAseq samples. “STAR” (https://github.com/alexdobin/STAR)^71^ was used as the aligner, and “Salmon” (https://github.com/COMBINE-lab/salmon)^72^ was used for quantification at both transcript and gene levels. Data quality was checked using different metrics, and the samples that met the following criteria were considered high-quality: sufficient sequencing depth (>20M reads), low duplication level (duplication < 70%), and good alignment (unique mapped rate > 70%). For this research, only gene-level quantification was considered, and two data formats were used. Read counts were used for differential expression (DE) analysis, while batch effect-corrected and normalized transcripts per million (TPM) values were used for gene expression and function-related analyses. Principal component analysis (PCA) was performed using the “prcomp” function in R. DE analysis was performed using the “DESeq2” package in the R environment (https://github.com/mikelove/DESeq2)^73^. DE genes were selected using the following cut-off conditions: adjusted p-value < 0.05, |log2 fold-change| > 0.5, and average count > 20.

The functional enrichment of DE genes was performed in two steps: 1. Gene-set enrichment analysis on selected up-regulated or down-regulated DE genes using Enrichr gene-set collections. This was performed using the “enrichR” R package (https://github.com/GuangchuangYu/enrichR)^33^. 2. Significantly enriched gene-sets were collected from the Enrichr website and used to perform single-sample gene-set enrichment (ssGSEA) on each transcriptome data in all three groups (control, CS-FBS, and ENZ). To visualize on a heatmap, the enrichment scores of each gene-set were scaled. The p-values were calculated using the student t-test.

For visualization, the heatmaps were generated using the “pHeatmap” R package (https://github.com/raivokolde/pheatmap). The volcano plot was generated by the “EnhancedVolcano” R package (https://github.com/kevinblighe/EnhancedVolcano). Bar plots were generated using the “ggplot2” R package (https://github.com/tidyverse/ggplot2)^74^.

### 3.4 ATACseq and ChIPseq data processing

The “nf-core/atacseq” pipeline was used for ATACseq data processing. The sequencing reads were aligned to GRCh38 genome using STAR^71^ with spliced alignment function disabled. The STAR was chosen over other aligners because of its faster speed. Then ATACseq peaks were called by MACS2^75^. The HOMER^76^ was used for peak annotation. The normalized bigwig files were generated using “bedGraphToBigWig” (https://genome.ucsc.edu/goldenpath/help/bigWig.html) and were used to generate heatmaps and profile plots for visualization. The GRCh38 genome annotiation file (.gtf) was loading into R environment and converted to 4 bed files containing the genomics regions of ADT-up, ADT-down, constant-high and constant-low genes.

ChIPseq data processing shared most of steps as ATACseq with only difference in peak calling. All ChIPseq samples were separated into 2 groups: chromatin modifications and transcription factors (co-factors) while the prior group of samples’ peak were called using “broad-peaks” setting and later samples were called using “narrow-peaks” given the fact that chromatin modifications are usually wide. For both ATACseq and ChIPseq, bigWig files were generated for visualization by deeptools, and peak files were produced for quantification and annotation.

We have deposited a total of 137 bigwig files and 468 peak-related files from ATACseq samples, along with 268 bigwig files and 604 peak-related files from ChIPseq samples on FigShare for future reuse. Some files were not released because of low quality. The individual accession link of these files were listed in the data availability section. Additionally, raw read counts, TPM values, and a list of differentially expressed genes from RNAseq samples have also been included. These bigwig and peak-related files are compatible with visualization tools like Integrative Genomics Viewer (IGV, https://igv.org/) or UCSC Genome Browser (https://genome.ucsc.edu/). The RNAseq data can be analyzed using R. Furthermore, we have developed a shinyApp named as LNCaP-ADT (https://pcatools.shinyapps.io/shinyADT/) where users can interactively explore gene expression profiles from RNAseq samples across control, CS-FBS, and ENZ groups.

## 4. Discussion

This study established a comprehensive multi-omics dataset for LNCaP cells, revealing insights into the impact of AR signaling on the transcriptome and epigenome under ADT conditions. AR signaling predominantly influences chromatin accessibility and modifications at distal enhancers, with AR binding observed at these enhancers rather than proximal promoters. Through this mechanism, AR induces the expression of genes related to prostatic differentiation and metabolism, while inhibiting those associated with neuron, inflammation, and Wnt signaling (Figure 4A). Notably, in LNCaP cells, genes induced by AR (inhibited by ADT) maintained a more active chromatin status compared to AR-inhibited genes, even under the ADT treatment. Furthermore, this study sheds light on NEPCa development, suggesting that ADT does not directly induce neuroendocrine differentiation in LNCaP cells. Additionally, we have made all this high-quality, reprocessed multi-omics data publicly available, providing a valuable resource for the research community to facilitate further studies and advancements in prostate cancer research (Figure 4B).

**Figure 4.**
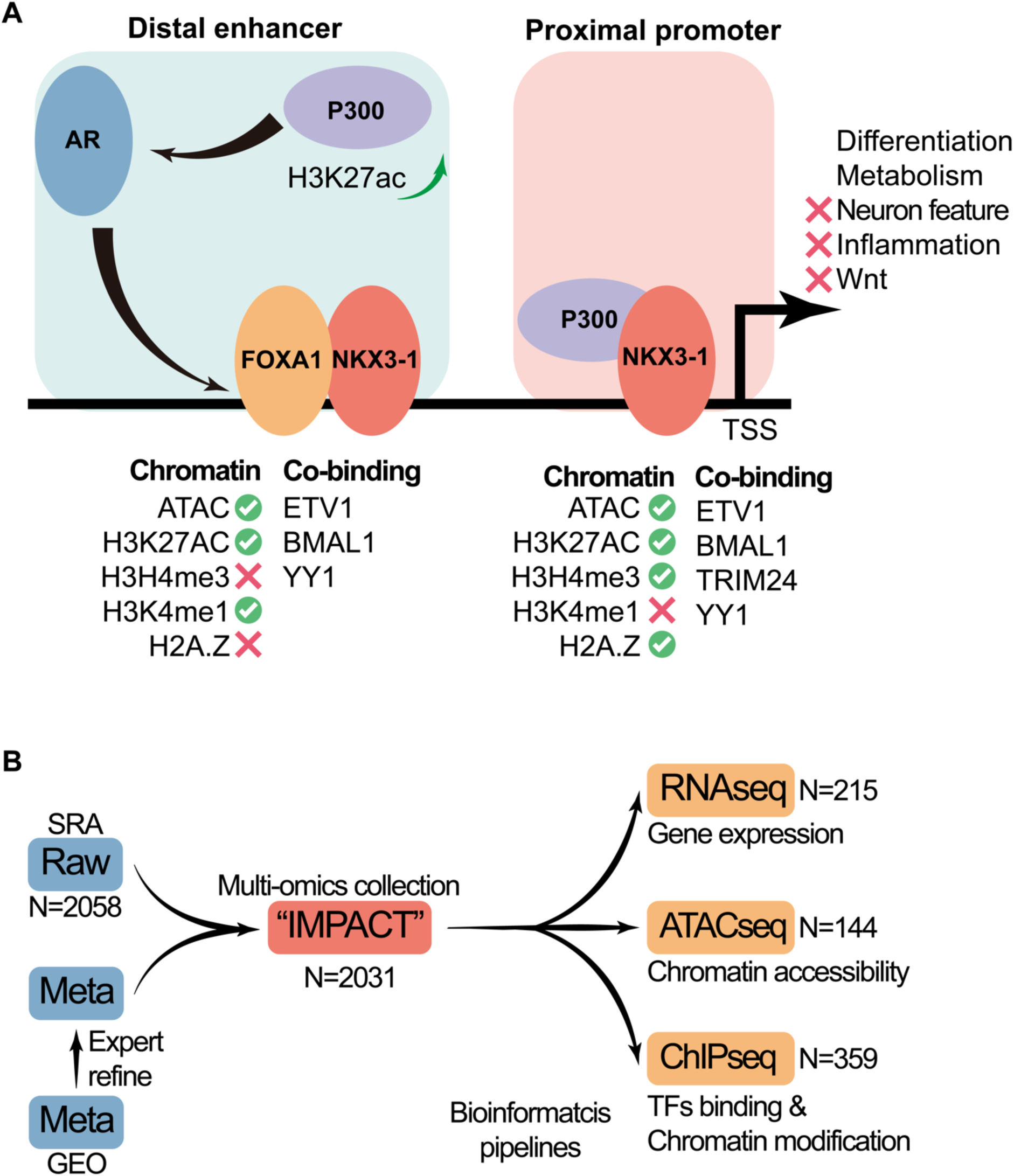
Summary Diagram. (A) An animation diagram illustrating the binding pattern and downstream function of AR. (B) The workflow of the present “IMPACT” research, starting from data acquisition to data redistribution.

Researchers worldwide are energetically and continually providing novel insights into cancer, including prostate cancer. A large amount of research has generated high-throughput data deposited in public cohorts, as indicated by the strikingly increasing number of samples. As shown in this research, thousands of multi-omics datasets can be retrieved from a single prostate cancer cell line. While these data have been interpreted by the original authors as contributions to the field, the high-dimensional nature of these high-throughput data means many hidden gems are still waiting to be discovered. More importantly, due to the limited resources of single research groups, it is almost impossible to describe the overall molecular properties, including the transcriptome and epigenome, of each cell line or cancer type through individual studies.

To address this, we propose the Integrative Multi-Omics Prostate Cancer Informatics Initiative (IMPACT), which reprocesses existing publicly available prostate cancer multi-omics data using state-of-the-art bioinformatics pipelines and reference genome. This study established the data collection and processing pipeline and the method for data redistribution. In future studies, more comprehensive analyses will be performed on other prostate cancer cell lines and clinical samples. We hope our bioinformatics studies can make prostate cancer research easier, resources more accessible, and the more unbiased.

## Acknowledgement

This research was supported by NIH R01 CA226285 to X. Yu, LSU Health Shreveport FWCC Carroll Feist postdoctoral Fellowship to S. Cheng, and LSU Health Shreveport FWCC Carroll Feist predoctoral Fellowship to L. Li. We would like to acknowledge the invaluable assistance of the Research IT team from LSUHS and extend special thanks to Sawyer Jarrod for the exceptional technical support with data sharing.

## Competing interests

The authors declare that they have no competing interests.

## Authors’ contributions

Conceptualization, Investigation, Methodology, Data curation, Software, Visualization: SC; Writing – original draft: LL, KC, SC; Writing – review & editing: XY, SC.

## Data availability

All data generated by this study were deposited for public access. For RNAseq samples, the read count, TPM and DE table files were deposited to https://doi.org/10.60688/lsuhs.26254769.v1. For ATACseq samples, the bigwig files were deposited to https://doi.org/10.60688/lsuhs.26253572.v1 (original) and https://doi.org/10.60688/lsuhs.26254280.v1 (scaled). The peak files were deposited to https://doi.org/10.60688/lsuhs.26254559.v1. For ChIPseq samples, the bigwig files were deposited to https://doi.org/10.60688/lsuhs.26262245.v1. The peak files were deposited to https://doi.org/10.60688/lsuhs.26264216.v1 and https://doi.org/10.60688/lsuhs.26264273.v1. In addition, an online application “LNCaP-ADT” was developed and launched publicly for users to access and visualize the data without coding requirement.

## Code availability

All codes generated for this research were deposited to https://github.com/schoo7/LNCaP-ADT.

## Declaration of generative AI and AI-assisted technologies in the writing process

During the preparation of this work the authors used ChatGPT in order to improve language and readability. After using this tool/service, the authors reviewed and edited the content as needed and take full responsibility for the content of the publication.

## Notes

https://doi.org/10.60688/lsuhs.26254769.v1

https://doi.org/10.60688/lsuhs.26253572.v1

https://doi.org/10.60688/lsuhs.26254280.v1

https://doi.org/10.60688/lsuhs.26254559.v1

https://doi.org/10.60688/lsuhs.26262245.v1

https://doi.org/10.60688/lsuhs.26264216.v1

https://doi.org/10.60688/lsuhs.26264273.v1

### Competing Interest Statement

The authors have declared no competing interest.

